# Free-water metrics in medial temporal lobe white matter tract projections relate to longitudinal cognitive decline

**DOI:** 10.1101/2020.01.06.896217

**Authors:** Derek B. Archer, Elizabeth E. Moore, Niranjana Shashikumar, Logan Dumitrescu, Kimberly R. Pechman, Bennett A. Landman, Katherine Gifford, Angela L. Jefferson, Timothy J. Hohman

## Abstract

**Objective:** Hippocampal volume is a sensitive marker of neurodegeneration and a well-established predictor of age-related cognitive impairment. Recently, free-water (FW) magnetic resonance imaging (MRI) has shown associations with pathology in Alzheimer’s disease (AD), but it is still unclear whether these metrics are associated with measures of cognitive impairment. Here, we investigate whether FW and FW-corrected fractional anisotropy (FA_T_) within medial temporal lobe white matter tracts (cingulum, fornix, uncinate fasciculus, inferior longitudinal fasciculus, and tapetum) provides meaningful contribution to cognition and cognitive decline beyond hippocampal volume.

**Participants and Methods:** Vanderbilt Memory & Aging Project participants (n=319, 73±7 years, 59% male) with normal cognition and mild cognitive impairment (40% of cohort) underwent baseline brain MRI, including structural MRI to quantify hippocampal volume, diffusion MRI to quantify medial temporal lobe white matter tract FW and FA_T_, and longitudinal neuropsychological assessment with a mean follow-up of 3.5 years. Linear regressions were conducted to determine how hippocampal volume and white matter tract FW and FA_T_ interact with baseline memory and executive function performances. Competitive model analyses determined the unique variance provided by white matter tract FW and FA_T_ beyond that of hippocampal volume and other comorbidities. Linear mixed-effects models were conducted to determine how baseline hippocampal volume and white matter tract FW and FA_T_ interact to explain longitudinal change in memory and executive function performances.

**Results:** FW in the inferior longitudinal fasciculus, tapetum, uncinate fasciculus, and cingulum were robustly associated with baseline memory and executive function. Further, competitive model analysis showed that tract FW contributed unique variance beyond other comorbidities and hippocampal volume for memory (ΔR_adj_^2^ range: 0.82-2.00%) and executive function (ΔR_adj_^2^ range: 0.88-1.87%). Longitudinal analyses demonstrated significant interactions of hippocampal volume and FA_T_ in the inferior longitudinal fasciculus (p=0.02), tapetum (p=0.02), uncinate fasciculus (p=0.02), and cingulum (p=0.002) with decline in memory. For decline in executive function, we found significant interactions of hippocampal volume and FA_T_ in inferior longitudinal fasciculus (p=0.03), tapetum (p=0.02), uncinate fasciculus (p=0.02), and fornix (p=0.02), as well as cingulum (p=0.02) and fornix (p=0.02) FW.

**Conclusions:** Our results highlight novel associations between FW and FA_T_ measures of medial temporal lobe tract microstructure and cognitive performance such that individuals with smaller hippocampal volumes and lower tract microstructure experience greater cognitive decline. These results suggest that white matter has a unique role in cognitive decline and, therefore, could be used to provide better disease staging, allowing for more precise disease monitoring in AD.

## Introduction

Alzheimer’s disease (AD) is a neurodegenerative disorder which is predominantly attributed to neurofibrillary tangles and β-amyloid plaques in the gray matter and is associated with cognitive impairment. As such, pathological biomarkers of AD which are associated with cognitive impairment and are predictive of cognitive decline are important for a more accurate assessment of disease severity and important for the development of novel therapies in AD. Neuroimaging biomarkers can provide an *in-vivo* assessment pathological biomarkers, and several gray matter specific biomarkers currently exist in AD. For example, structural magnetic resonance imaging (MRI) can quantify hippocampal atrophy (Thompson *et al*., 2004; Apostolova *et al*., 2006; Apostolova *et al*., 2010) and positron emission tomography can detect deposition of amyloid and tau in the cingulate cortex, middle temporal gyrus, and frontal cortex (Klunk *et al*., 2004; Schwarz *et al*., 2016). Since AD begins primarily in the gray matter, such as the medial temporal lobe, it is logical that gray matter biomarkers would be at the forefront of AD research; however, it is well-known that AD is a network level disease that spreads from the medial temporal lobe to remote portions of the brain via white matter tracts. Accordingly, post-mortem studies of AD have suggested that neuronal loss and synaptic pathology within the hippocampus and frontal cortex are also associated with cognitive impairment (DeKosky and Scheff, 1990; Andrade-Moraes *et al*., 2013), supporting the idea that AD propagates from the medial temporal lobe to frontal lobe via white matter tracts transneuronally (Caso *et al*., 2015). Further, recent studies of white matter microstructure in preclinical AD have shown robust associations with CSF measures of neurofibrillary tangle load (CSF p-tau), thin unmyelinated axons (CSF t-tau), and soluble amyloid precursor protein beta (sAPPβ) (Hoy *et al*., 2017). While these studies shed light into the relationship between white matter microstructure and AD pathology, it is still unclear what relationship white matter microstructure may have on cognitive decline in AD. It is also unknown whether measures of gray matter and white matter have a synergistic effect on cognitive decline. Therefore, this study sought to determine the association of white matter tract microstructure with baseline and future cognitive impairment in a large longitudinal cohort of individuals with varying levels of hippocampal atrophy.

Diffusion MRI (dMRI) is a particularly useful neuroimaging technique as it allows for the *in-vivo* quantification of white matter tract microstructural deficits. The most well-established microstructural measure in dMRI is fractional anisotropy (FA), and FA reductions have been associated with demyelination and axonal degradation (Beaulieu, 2002). With respect to AD, the importance of white matter tract microstructure, particularly in the fornix and cingulum bundle, have been previously described. For example, lower fornix FA has been found in individuals with presenilin-1 and presenilin-2 genetic mutations for familial AD, and lower FA was a stronger predictor of cognitive deficits than hippocampal volume (Mielke *et al*., 2012). Lower cingulum bundle FA has also been reported in AD compared to individuals with normal cognition or with amnestic mild cognitive impairment (MCI), and microstructure of the cingulum bundle was associated with global cognition (Bozzali *et al*., 2012). FA in the uncinate fasciculus has also been shown to be lower in AD (Yasmin *et al*., 2008). While these finding have suggested a significant role of white matter in AD, FA is susceptible to partial volume effects (i.e., both fluid and tissue are present within a voxel) and could therefore be limiting our findings in AD. Fortunately, new techniques, such as free-water (FW) imaging, have allowed for the separation of the fluid (FW) and tissue (FW-corrected FA [FA_T_]) components, thus increasing the biological specificity of dMRI studies. To date, no study has comprehensively quantified medial temporal lobe white matter FW and FA_T_ and determined how these measures are associated with cognitive impairment and cognitive decline.

The present study conducted high-resolution probablistic tractography of the medial temporal lobe projections, including the cingulum bundle, inferior longitudinal fasciculus, and uncinate fasciculus, in conjunction with freely available white matter tract templates of the fornix (Brown *et al*., 2017) and tapetum (Archer *et al*., 2019), to determine unique contributions of white matter FW and FA_T_ in age-related cognitive impairment in a longitudinal cohort of individuals with normal cognition and MCI. Furthermore, we examined the synergistic relationship of gray matter atrophy (i.e., hippocampal volume) and medial temporal lobe white matter FW and FA_T_ in age-related cognitive decline. Our hypothesis was that medial temporal lobe tract microstructure would explain unique variance in baseline cognitive performance and that individuals with both reduced medial temporal lobe tract microstructure and smaller hippocampal volume would have a more rapid rate of cognitive decline over the follow-up period.

## Methods

### Study Cohort

The Vanderbilt Memory & Aging Project was launched in 2012 and is a longitudinal observational study. Cohort inclusion criteria required participants to be 60 years of age or older, speak English, have adequate auditory and visual acuity for testing, and have a reliable study partner. Full characterization of cohort has been described elsewhere (Jefferson *et al*., 2016). At study entry, participants completed a comprehensive neuropsychological examination and were placed into three categories – cognitively normal, early mild cognitive impairment (eMCI), and MCI (Kresge *et al*., 2018). Neuropsychological assessment was collected longitudinally up to five years. Apolipoprotein E (*APOE*) haplotype status (ε2, ε3, ε4) was determined using single-nucleotide polymorphisms (SNPs) rs7412 and rs429358, which were genotyped using TaqMan on frozen whole blood. T1-weighted MRI and dMRI was also collected (see acquisition and preprocessing of MRI data below). The protocol was approved by the Vanderbilt University Medical Center Institutional Review Board, and written informed consent was obtained prior to data collection.

### Neuropsychological Composites

The neuropsychological protocol was performed by experienced technicians who assessed several cognitive domains, including episodic memory and executive function. As previously described (Kresge *et al*., 2018), an executive function composite was calculated using the Delis-Kaplan Executive Function System (DKEFS) Tower Test, DKEFS Letter-Number Switching, DKEFS Color-Word Inhibition, and Letter Fluency (FAS). For the memory composite, the California Verbal Learning Test-Second Edition (CVLT-II) Total Learning, Interference Condition, Long Delay Free Recall, and Recognition components were used in addition to the identical components of the Biber Figure Learning Test. Composites were created using latent variable models, and the final composites represent z scores.

### Acquisition and Quantification of Hippocampal Volume and Hippocampal Atrophy

T1-weighted MRI images (TR/TE: 8.9/4.6 ms, resolution: 1mm isotropic) were collected from each participant on a 3T Philips Achieva system (Best, The Netherlands) using an 8-channel SENSE reception coil. Multi-Atlas segmentation was conducted to obtain hippocampal segmentations and volumes (Asman and Landman, 2012). Hippocampal volume was quantified by averaging the left and right hippocampal masks and then standardized by total intracranial volume. Based on a prior study, participants were classified into two groups based on hippocampal volume – neurodegenerative negative (>6723 mm^3^) and neurodegenerative positive (<6723 mm^3^) (Mormino *et al*., 2014).

### dMRI Acquisition and Preprocessing

dMRI images (TR/TE: 10,000/60 ms, resolution: 2mm isotropic, b-values: 0, 1,000 s/mm^2^) were collected from each participant using the same scanner described above. Images were collected along 32 diffusion gradient vectors and five non-diffusion weighted images (B_0_). FSL 6.0.1 (fsl.fmrib.ox.ax.uk) was used for all dMRI preprocessing (Jenkinson *et al*., 2012; Andersson and Sotiropoulos, 2016). Quality assessment of all dMRI scans was performed manually. The data were first corrected for head motion using an affine registration and the brain was then extracted from the skull. This corrected image was used as input in two different procedures: (1) DTIFIT to calculate fractional anisotropy maps (FA), and (2) custom written MATLAB (R2019a, The Mathworks, Natick, MA, USA) code to calculate FW and FA_T_ maps, as described in detail previously (Pasternak *et al*., 2009; Archer *et al*., 2019). To obtain a standardized space representation of the FW and FA_T_ maps, the FA_T_ map was registered to an in-house FA_T_ template (1mm isotropic) by a nonlinear warp using the Advanced Normalization Tools (ANTs) package (Avants *et al*., 2008). This nonlinear warp was applied to the FW map.

### White Matter Tractography Templates

Human Connectome Project (HCP) data was used to create white matter tract templates using an established approach (Archer *et al*., 2018). White matter tract templates included tracts projecting from hippocampus (tapetum, fornix, cingulum bundle, uncinate fasciculus, and inferior longitudinal fasciculus). For the tapetum, we used a template tract from the Transcallosal Tract Template (TCATT) (Archer *et al*., 2019). For the fornix, we used a well-established fornix template (Brown *et al*., 2017). For the cingulum bundle, uncinate fasciculus, and inferior longitudinal fasciculus, we conducted probabilistic tractography to create white matter tract templates as current tractography templates for these tracts were either incomplete or nonexistent. Consistent with prior work (Archer *et al*., 2018; Archer *et al*., 2019), we conducted probabilistic tractography using the probtrackx2 program in FSL using default settings (samples: 5,000, curvature threshold: 0.2, FA threshold: 0.2) on 100 individuals from the HCP (Van Essen *et al*., 2013). For the cingulum bundle, we used a hippocampal mask as a seed and an anterior cingulate cortex mask as a waypoint. For the uncinate fasciculus, we used a hippocampal mask as a seed and a prefrontal cortex mask as a waypoint. For the inferior longitudinal fasciculus, we used a hippocampal mask as a seed and a posterior parietal cortex mask as a waypoint. FA_T_ and FW were then calculated within all tracts for all participants.

### Statistical Analyses

All statistical analyses were performed in R version 3.5.2 (http://www.r-project.org/). Covariates included age, sex, education, cognitive status, race/ethnicity, Framingham Stroke Risk Profile (FSRP) scores (Wolf *et al*., 1991; D’Agostino *et al*., 1994), and *APOE*-ε4 carrier status. *APOE*-ε4 carrier status was defined as positive (ε2/ε4, ε3/ ε4, ε4/ ε4) or negative (ε2/ε2, ε2/ε3, ε3/ε3). Significance was set a priori as α=0.05 and correction for multiple corrections were made using the false discovery rate (FDR) method.

For all analyses, only one tract per model was considered. Baseline effects of all tracts (tapetum, fornix, cingulum, uncinate fasciculus, and inferior longitudinal fasciculus) for both measures (FW and FA_T_) were estimated using a general linear model for each of the 3 outcomes (hippocampal volume, memory composite, executive function composite). Further, *cognitive status x white matter tract* interaction terms (e.g., cognitive status x cingulum FA_T_) were evaluated for each tract and measure on the 3 outcomes. We then conducted an analysis of *hippocampal volume x white matter tract* interaction terms on memory and executive function composites. All significant terms were followed with post-hoc competitive model analysis which assessed the unique variance white matter measures contributed to cognitive function beyond comorbities and hippocampal volume.

Finally, we evaluated the interaction between baseline hippocampal volume and white matter tract FW and FA_T_ on longitudinal memory and executive function composites using mixed-effects regression. Time was modeled as years from baseline for each participant, and both time and the intercept were inputted as both fixed and random effects in the model. The three-way *hippocampal volume x white matter tract x time* term assessed baseline interaction effects on longitudinal change in cognition. All lower-order interactions were included in the model.

## Results

### Participant Characteristics

Demographic and clinical variables for each cognitive status group (cognitively normal, eMCI, MCI) are summarized in **Table 1**. Patients were mostly well-educated, elderly, non-Hispanic white individuals. There were no significant differences in age, sex distribution, or race distribution between groups. The cognitively normal group had more education than the MCI group. The MCI group had more APOE4 positive individuals than the cognitively normal group. As expected, the there were significant differences between the cognitively normal, eMCI, and MCI groups in hippocampal volume, total FSRP score, and neuropsychological composites.

**Table 1 –.**
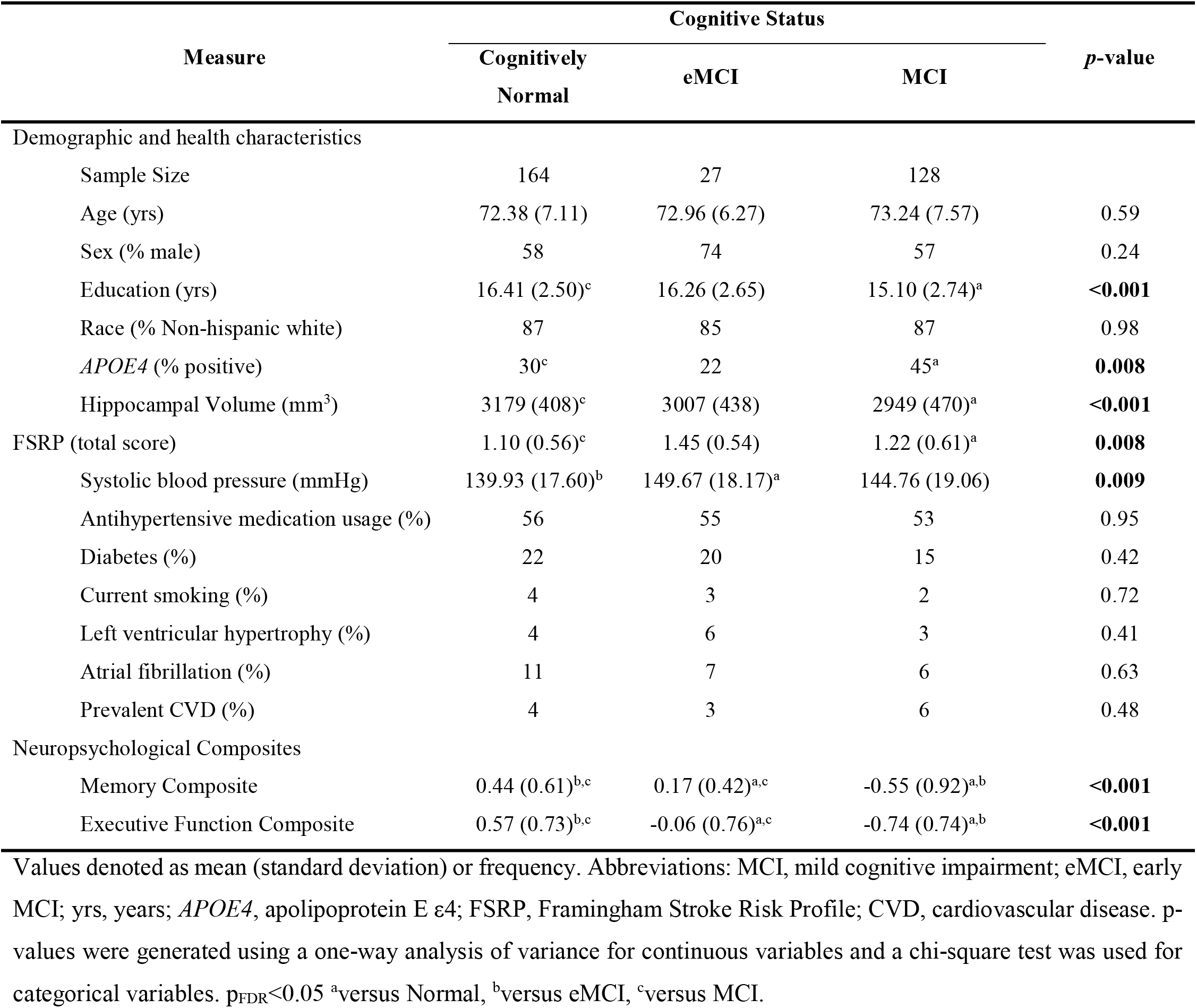
Demographic and Health Characteristics

| Measure | Cognitive Status |  |  | *p*-value |
| --- | --- | --- | --- | --- |
|  | Cognitively Normal | eMCI | MCI |  |
| Demographic and health characteristics |  |  |  |  |
| Sample Size | 164 | 27 | 128 |  |
| Age (yrs) | 72.38 (7.11) | 72.96 (6.27) | 73.24 (7.57) | 0.59 |
| Sex (% male) | 58 | 74 | 57 | 0.24 |
| Education (yrs) | 16.41 (2.50)<sup>c</sup> | 16.26 (2.65) | 15.10 (2.74)<sup>a</sup> | **<0.001** |
| Race (% Non-hispanic white) | 87 | 85 | 87 | 0.98 |
| *APOE4* (% positive) | 30<sup>c</sup> | 22 | 45<sup>a</sup> | **0.008** |
| Hippocampal Volume (mm<sup>3</sup>) | 3179 (408)<sup>c</sup> | 3007 (438) | 2949 (470)<sup>a</sup> | **<0.001** |
| FSRP (total score) | 1.10 (0.56)<sup>c</sup> | 1.45 (0.54) | 1.22 (0.61)<sup>a</sup> | **0.008** |
| Systolic blood pressure (mmHg) | 139.93 (17.60)<sup>b</sup> | 149.67 (18.17)<sup>a</sup> | 144.76 (19.06) | **0.009** |
| Antihypertensive medication usage (%) | 56 | 55 | 53 | 0.95 |
| Diabetes (%) | 22 | 20 | 15 | 0.42 |
| Current smoking (%) | 4 | 3 | 2 | 0.72 |
| Left ventricular hypertrophy (%) | 4 | 6 | 3 | 0.41 |
| Atrial fibrillation (%) | 11 | 7 | 6 | 0.63 |
| Prevalent CVD (%) | 4 | 3 | 6 | 0.48 |
| Neuropsychological Composites |  |  |  |  |
| Memory Composite | 0.44 (0.61)<sup>b,c</sup> | 0.17 (0.42)<sup>a,c</sup> | -0.55 (0.92)<sup>a,b</sup> | **<0.001** |
| Executive Function Composite | 0.57 (0.73)<sup>b,c</sup> | -0.06 (0.76)<sup>a,c</sup> | -0.74 (0.74)<sup>a,b</sup> | **<0.001** |

Values denoted as mean (standard deviation) or frequency. Abbreviations: MCI, mild cognitive impairment; eMCI, early MCI; yrs, years; *APOE4*, apolipoprotein E ε4; FSRP, Framingham Stroke Risk Profile; CVD, cardiovascular disease. *p*-values were generated using a one-way analysis of variance for continuous variables and a chi-square test was used for categorical variables. $p_{FDR} < 0.05$ <sup>a</sup>versus Normal, <sup>b</sup>versus eMCI, <sup>c</sup>versus MCI.

### Baseline Tract Microstructure Association with Baseline Hippocampal Volume and Memory/Executive Composites

Baseline results are presented in **Table 2** and graphically summarized in **Figure 1**. White matter associations with hippocampal volume included FA_T_ in the tapetum, uncinate fasciculus, cingulum, and fornix (all p≤0.02) and FW in the inferior longitudinal fasciculus, uncinate fasciculus, cingulum, and fornix (all p<0.009). White matter associations with memory and executive function included FW in the inferior longitudinal fasciculus, tapetum, uncinate fasciculus, and cingulum (all p≤0.04), but no FA_T_ associations with baseline cognitive performance were observed. As shown in **Figure 1A**, higher FW in the inferior longitudinal fasciculus was associated with lower composite memory performance in all cognitive status groups. Similarly, **Figure 1B** shows that higher FW in the tapetum is associated with lower composite executive performance in all cognitive status groups. Sensitivity analyses showed that significant associations did not differ by baseline cognitive status group (all p=0.94).

**Table 2 –.** Baseline Tract Microstructure Association with Baseline Hippocampal Volume, Composite Memory Performance, and Composite Executive Function Performance

| Tract | FA<sub>T</sub> |  | FW |  |
| --- | --- | --- | --- | --- |
|  | β (95% CI) | *p*-value | β (95% CI) | *p*-value |
| **Association with Baseline Hippocampal Volume** |  |  |  |  |
| Inferior Longitudinal Fasciculus | 2180 (1039 to 3322) | 0.06 | -3403 (-4070 to -2735) | **<0.001** |
| Tapetum | 3711 (2213 to 5210) | **0.02** | -1102 (-1755 to -448) | 0.10 |
| Uncinate Fasciculus | 3559 (2259 to 4858) | **0.01** | -2850 (-3456 to -2245) | **<0.001** |
| Cingulum Bundle | 3431 (2167 to 4694) | **0.01** | -3813 (-4553 to -3073) | **<0.001** |
| Fornix | 3239 (1909 to 4569) | **0.02** | -988 (-1342 to -634) | **0.008** |
| **Association with Baseline Memory Composite** |  |  |  |  |
| Inferior Longitudinal Fasciculus | 2.52 (0.54 to 4.5) | 0.37 | -4.57 (-5.78 to -3.35) | **0.01** |
| Tapetum | -1.19 (-3.81 to 1.43) | 0.81 | -2.79 (-3.91 to -1.67) | **0.04** |
| Uncinate Fasciculus | -0.61 (-2.9 to 1.67) | 0.83 | -2.85 (-3.96 to -1.75) | **0.04** |
| Cingulum Bundle | 1.29 (-0.93 to 3.51) | 0.75 | -3.73 (-5.09 to -2.36) | **0.03** |
| Fornix | 2.47 (0.14 to 4.79) | 0.41 | -0.97 (-1.59 to -0.35) | 0.24 |
| **Association with Baseline Executive Function Composite** |  |  |  |  |
| Inferior Longitudinal Fasciculus | 2.33 (0.39 to 4.26) | 0.38 | -2.93 (-4.13 to -1.73) | **0.04** |
| Tapetum | 4.29 (1.74 to 6.84) | 0.21 | -3.71 (-4.79 to -2.63) | **0.01** |
| Uncinate Fasciculus | 0.8 (-1.43 to 3.03) | 0.81 | -3.25 (-4.32 to -2.17) | **0.02** |
| Cingulum Bundle | 2.44 (0.27 to 4.61) | 0.40 | -3.43 (-4.76 to -2.10) | **0.04** |
| Fornix | 0.2 (-2.07 to 2.47) | 0.93 | -0.24 (-0.85 to 0.36) | 0.81 |

Abbreviations: FA<sub>T</sub>, free-water corrected fractional anisotropy; FW, free-water. Boldface signifies $p < 0.05$ .

**Figure 1 –.**
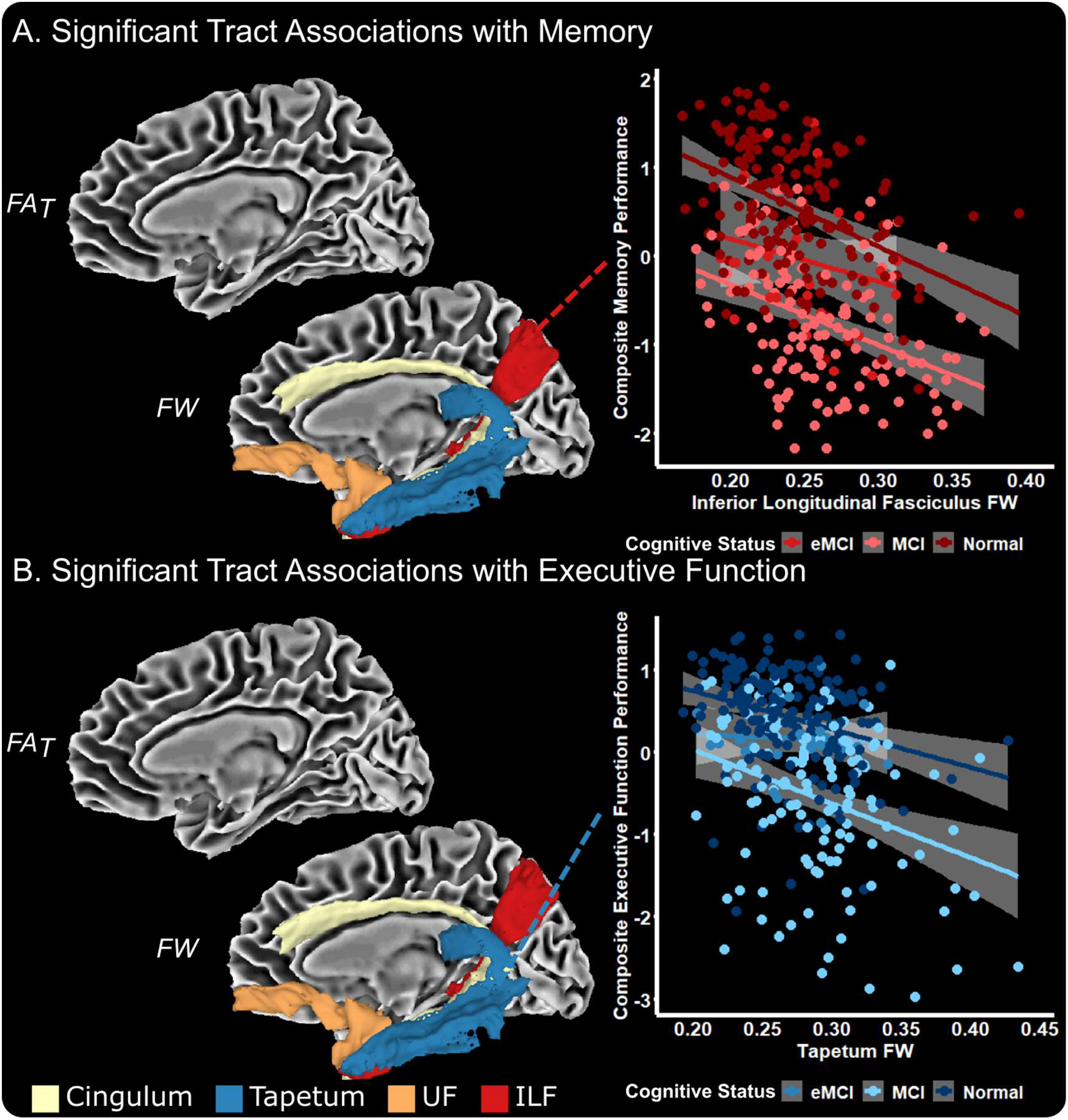
Baseline Associations with Composite Memory and Executive Function Performance. The medial temporal lobe tract measures which were associated with memory (**A**) and executive function (**B**) performance include FW in the cingulum bundle, tapetum, uncinate fasciculus (UF), and inferior longitudinal fasciculus (ILF). The association of ILF FW with memory performance (**A**) and tapetum with executive function performance (**B**) is shown.

We then conducted a competitive model analysis to determine the unique variance explained by hippocampal volume and white matter tract microstructure in addition to all covariates (age, sex, education, cognitive status, APOE-ε4 status, race, FSRP scores) (**Table 3**). We found that using only covariates there was a strong relationship with baseline memory (R_adj_^2^=51.02%; p=2.2×10^−16^). When adding hippocampal volume to this model, we found a small increase (ΔR_adj_^2^=0.58%) in the overall model (R_adj_^2^=51.60%; p=2.2×10^−16^). We then iteratively added tract FW and FA_T_ measures to this model to determine if tract microstructure explained any unique variance beyond covariates and hippocampal volume. We found that tract FA_T_ did not significantly contribute to the models; however, for FW, all tracts (aside from the fornix) were significant contributors to the model and provided increases in the R_adj_^2^ (range: 0.82%-2.00%). For executive function performance, there was a strong relationship with covariates along (R_adj_^2^=45.52%; p=2.2×10^−16^), and the addition of hippocampal volume to this model provided a reduction (ΔR_adj_^2^=-0.64%) in the overall model (R_adj_^2^=44.88%; p=2.2×10^−16^). We then added tract FW and FA_T_ measures to this model to determine if tract microstructure explained any unique variance beyond covariates and hippocampal volume to executive function performance. For FA_T_, we found that no tracts were significant contributors to the model. For FW, we found that all tracts (aside from the fornix) were significant contributors to the model and provided increases in the R_adj_^2^ (range: 0.88%-1.87%).

**Table 3 –.** Baseline Tract Microstructure x Hippocampal Interaction Associations

|  | Memory |  |  |  | Executive Function |  |  |  |
| --- | --- | --- | --- | --- | --- | --- | --- | --- |
| | $\beta$ | SE | *p*-value | $\Delta R_{adj}^2$ | $\beta$ | SE | *p*-value | $\Delta R_{adj}^2$ |
| Covariates + | (R<sub>adj</sub><sup>2</sup>=51.02%; p=2.2x10<sup>-16</sup>) |  |  |  | (R<sub>adj</sub><sup>2</sup>=45.52%; p=2.2x10<sup>-16</sup>) |  |  |  |
| Hippocampal Volume | 2.55x10<sup>-4</sup> | 9.80x10<sup>-5</sup> | **0.01** | 0.58 | 1.23x10<sup>-4</sup> | 9.60x10<sup>-5</sup> | 0.20 | -0.64 |
| Covariates + Hippocampal Volume + | (R<sub>adj</sub><sup>2</sup>=51.60%; p=2.2x10<sup>-16</sup>) |  |  |  | (R<sub>adj</sub><sup>2</sup>=44.88%; p=2.2x10<sup>-16</sup>) |  |  |  |
| ILF FA<sub>T</sub> | 2.52 | 1.97 | 0.20 | 0.10 | 2.33 | 1.93 | 0.23 | 0.08 |
| Tapetum FA<sub>T</sub> | -1.91 | 2.63 | 0.65 | -0.12 | 4.29 | 2.55 | 0.09 | 0.33 |
| UF FA<sub>T</sub> | -0.614 | 2.29 | 0.79 | -0.14 | 0.80 | 2.23 | 0.72 | -0.16 |
| Cingulum Bundle FA<sub>T</sub> | 1.29 | 2.22 | 0.56 | -0.10 | 2.44 | 2.17 | 0.26 | 0.05 |
| Fornix FA<sub>T</sub> | 2.47 | 2.32 | 0.29 | 0.02 | 0.20 | 2.27 | 0.93 | -0.18 |
| ILF FW | -4.57 | 1.22 | **<0.001** | 2.00 | -2.93 | 1.20 | **0.02** | 0.88 |
| Tapetum FW | -2.79 | 1.12 | **0.01** | 0.82 | -3.71 | 1.09 | **<0.001** | 1.87 |
| UF FW | -2.85 | 1.10 | **0.01** | 0.89 | -3.25 | 1.07 | **0.003** | 1.44 |
| Cingulum Bundle FW | -3.73 | 1.36 | **0.01** | 1.01 | -3.43 | 1.33 | **0.01** | 1.00 |
| Fornix FW | -0.97 | 0.62 | 0.12 | 0.23 | -0.24 | 0.61 | 0.69 | -0.15 |

Abbreviations: FA<sub>T</sub>, free-water corrected fractional anisotropy; FW, free-water; ILF, inferior longitudinal fasciculus; UF, uncinate fasciculus. B, SE, and *p*-values represent the parameter estimate for each variable. Boldface signifies *p*<0.05.

We then determined if there were baseline hippocampal x white matter tract interactions on memory and executive performance. The results are presented in **Supplemental Table 1**, which shows there were no significant interactions between white matter tract FA_T_ or FW and hippocampal volume on baseline memory or executive function performance.

### Tract-by-hippocampal Volume Interaction on Longitudinal Memory/Executive Composite

For annual change in memory performance, we found significant interactions between hippocampal volume and FA_T_ in the inferior longitudinal fasciculus, tapetum, uncinate fasciculus, and cingulum (all p≤0.02). No significant interactions were found with FW. For annual change in executive function performance, we found significant interactions between hippocampal volume and FA_T_ in the inferior longitudinal fasciculus, tapetum, uncinate fasciculus, and fornix (all p≤0.04), and with FW in the cingulum and fornix (all p≤0.02). Results for all models can be found in **Table 4**. **Figure 2** provides a summary of our findings for the annual change in memory and executive function. As shown in **Figure 2A**, lower FA_T_ in the cingulum is associated with a more rapid decline in memory performance, particularly among individuals with baseline neurodegeneration in the hippocampus. **Figure 2B** shows that higher FW in the fornix is associated with a more rapid decline in executive function, particularly among individuals with neurodegeneration in the hippocampus.

**Table 4 –.**
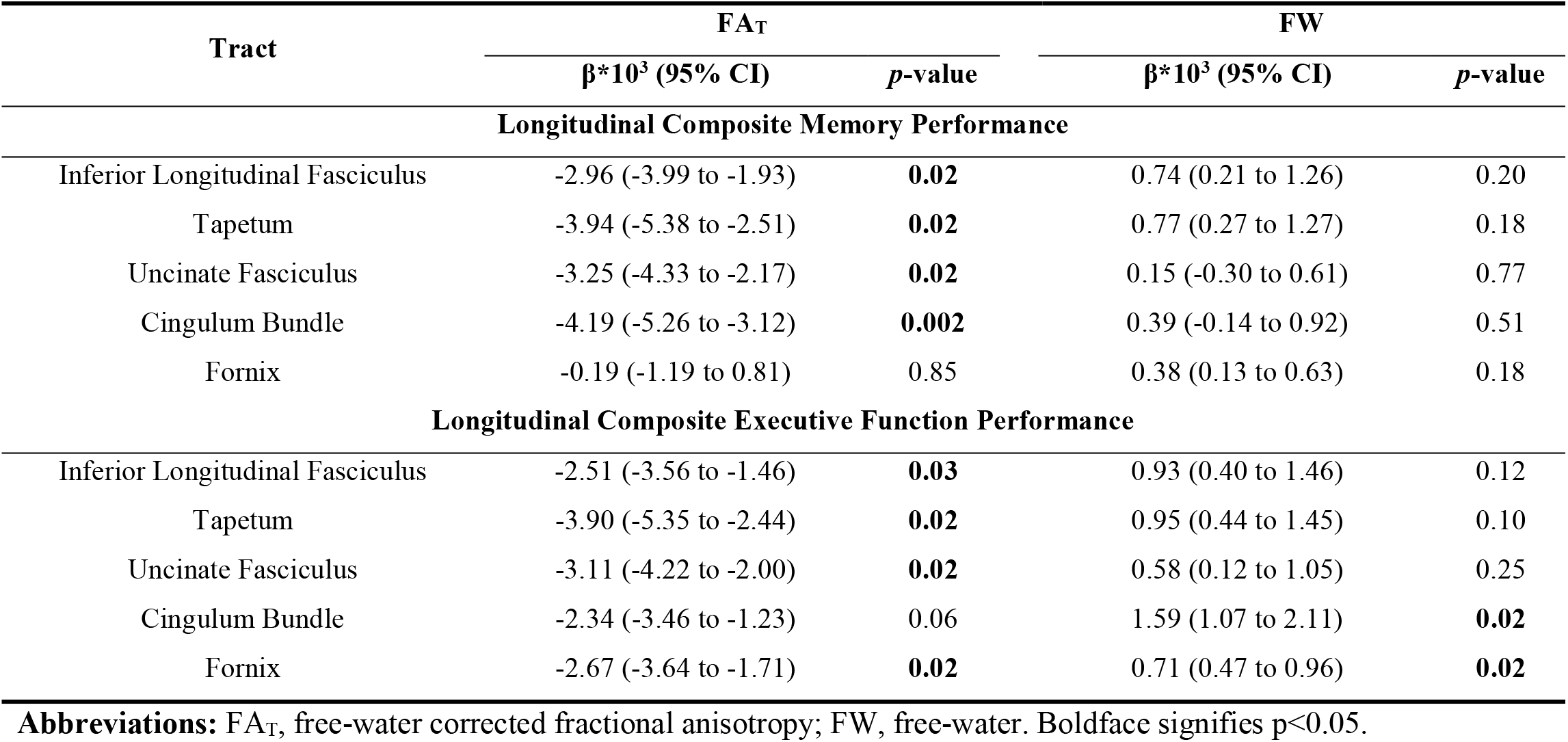
Baseline Tract Microstructure x Hippocampal Interaction Associations

**Figure 2 –.**
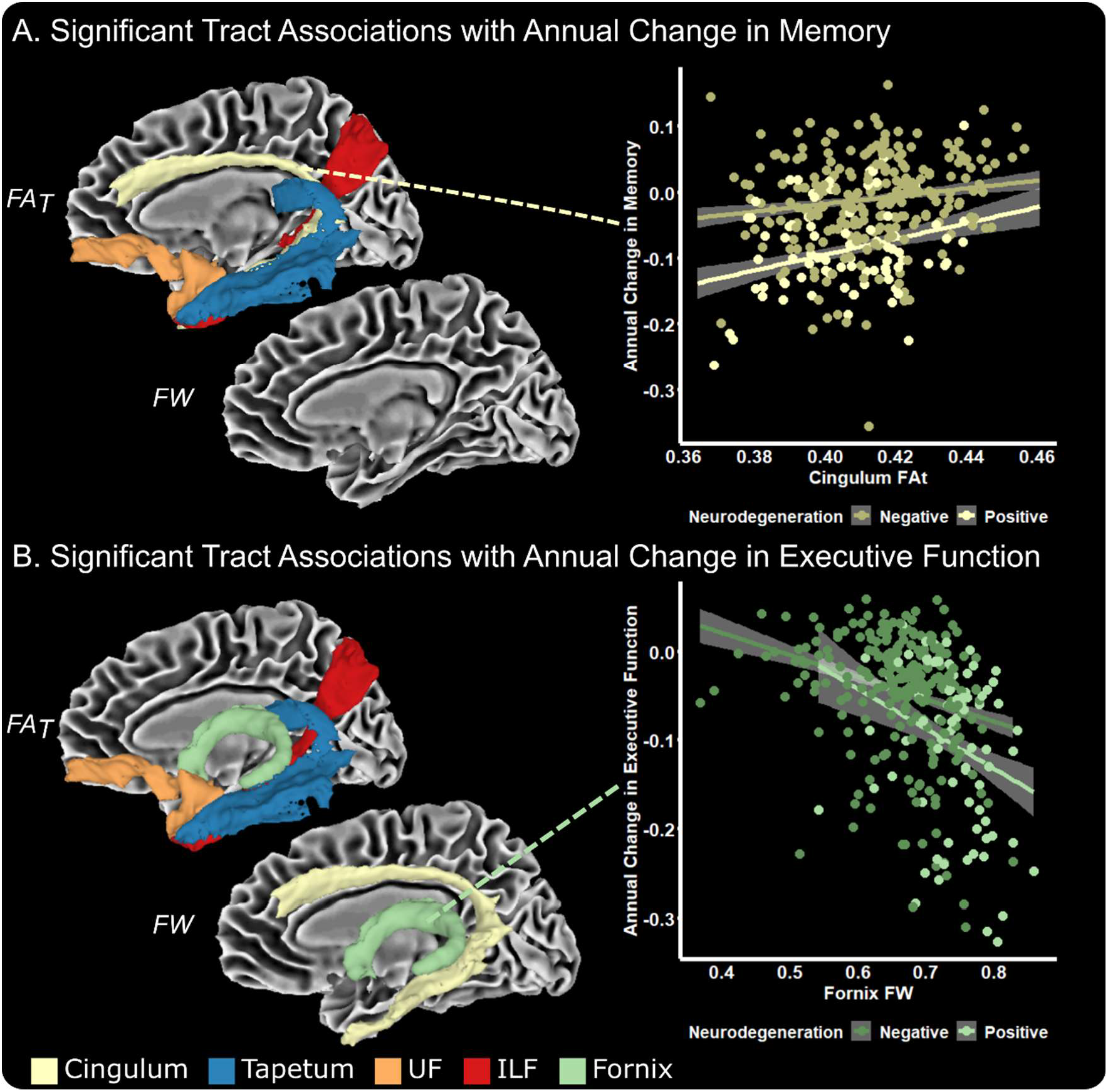
Baseline Tract x Hippocampal Interaction on Annual Change in Memory and Executive Function. The medial temporal lobe tract measures which had a significant interaction with hippocampal volume for annual change in memory include FA_T_ in the uncinate fasciculus (UF), inferior longitudinal fasciculus (ILF), cingulum bundle, and tapetum (**A**). For annual change in executive function performance, temporal tract measures which had a significant interaction with hippocampal volume include FA_T_ in the UF, ILF, cingulum bundle, and tapetum, as well as FW in the UF, ILF, tapetum, and fornix. Hippocampal neurodegeneration groups (negative, positive) are based on a previously identified cutoff value for hippocampal neurodegeneration (positive: volume ≤ 6723 mm^3^) (Mormino *et al*., 2014).

## Discussion

The present study examined relationships between hippocampal volume and the microstructure of the medial temporal lobe white matter tracts to evaluate how these sensitive imaging metrics relate to cognitive performance and cognitive decline. Specifically, we used freewater imaging, an innovative technique which overcomes the limitations of conventional dMRI, to quantify microstructural values (FW and FA_T_) in several medial temporal lobe projections, including the cingulum bundle, fornix, tapetum, inferior longitudinal fasciculus, and uncinate fasciculus. We report three main findings. First, we found that FW and FA_T_ in medial temporal lobe tracts were strongly associated with hippocampal volume. Second, we found that baseline measures of medial temporal lobe white matter tract FW were robustly associated with baseline cognitive performance. Competitive model analyses determined that FW in the tapetum, cingulum, uncinate fasciculus, and inferior longitudinal fasciculus explained variance above and beyond hippocampal volume and covariates in cognitive performance. Third, we found significant interactions of hippocampal volume and medial temporal lobe white matter tract FA_T_ on longitudinal cognitive trajectory, whereby individuals with lower hippocampal volume and lower white matter tract FA_T_ exhibited greater longitudinal decline. This study therefore provides direct evidence that medial temporal lobe white matter projections are relevant to cognitive performance even when statistically accounting for hippocampal atrophy.

The medial temporal lobe white matter tract microstructure association with memory and executive function performance is consistent with previous reports (Mielke *et al*., 2012; Ji *et al*., 2019); however, the mechanisms by which medial temporal lobe microstructure is linked to memory and executive function is unclear. One prevailing hypothesis is that white matter microstructure is deficient in AD as a result of Wallerian-degeneration following hippocampal atrophy (Sachdev *et al*., 2013). Alternatively, contrasting research has suggested that white matter degeneration may precede noticeable hippocampal atrophy (Zhuang *et al*., 2013; Hoy *et al*., 2017; Metzler-Baddeley *et al*., 2019). While these hypotheses differ on the temporal ordering of white matter decline, it is clear that white matter measures are associated with cognition (Bozzali *et al*., 2012; Mielke *et al*., 2012). Our results add to this growing body of literature by demonstrating that hippocampal volume synergistically interacts with white matter microstructure to explain longitudinal decline in memory and executive function and highlights the unique contribution of white matter changes in explaining cognitive decline over the course of aging and disease. Future longitudinal studies should determine if white matter decline in AD is an independent process, a consequence of hippocampal atrophy, or a combination of these processes.

One interesting observation is that while medial temporal lobe white matter FW is robustly associated with baseline memory and executive function performances, medial temporal lobe FA_T_ interacts with hippocampal volume to explain longitudinal decline in memory and executive function. These two white matter metrics are thought to reflect different neurobiological processes. FW measures unbound water molecules in the white matter, and thus higher FW could reflect a neuroinflammatory or more general axonal damage process (Pasternak *et al*., 2012). Therefore, the finding that higher FW is associated with lower baseline cognitive performance may reflect a more advanced neurodegenerative state in which atrophy has already impacted the white matter. In contrast, FA_T_ is a more direct measure of white matter microstructure, as it calculates intracellular white matter microstructure, with lower FA_T_ indicating more white matter vulnerability. Thus, while we found that FA_T_ does not appear to be as sensitive to baseline cognitive performance as FW, our findings indicate that individuals with higher white matter vulnerability (i.e., lower FA_T_) are predisposed to a more rapid disease progression.

Accordingly, white matter vulnerability in AD has been directly linked to amyloid and tau pathology. An *in vitro* study of hippocampal neurons found that Aβ1-42 induced axonal degeneration that preceded cell death (Alobuia *et al*., 2013). Further, a biochemical analysis of AD white matter found that higher Aβ_40_ and Aβ_42_ was accompanied with lower myelin basic protein and ultimately demyelination (Roher *et al*., 2002). More recently, a tractography study using FW imaging in preclinical familial AD found that FW measures were significantly associated with both CSF amyloid and tau pathology (Hoy *et al*., 2017). Another study in familial AD showed that diffusion alterations in white matter tracts were associated with lower CSF Aβ_1-42_, but higher total tau, hyperphosphorylated tau (P-tau_181_), and microglial activation (Araque Caballero *et al*., 2018). It is therefore possible that our findings linking lower white matter FA_T_ to more rapid cognitive decline could be a direct result of AD pathology or could reflect an independent disease process of white matter injury that results in an enhanced susceptibility to cognitive decline. It will be critical for future research to better understand the mechanistic contributors to the observed FA_T_ effects and clarify where they fall within well-established neurodegenerative cascades.

The present study has several strengths, including a well-characterized longitudinal cohort and the application of free-water imaging, which overcomes the limitations of conventional dMRI techniques. An additional strength of this study is the incorporation of novel medial temporal lobe white matter tract templates. The use of white matter tract templates increases consistency between studies as the identical voxels are being evaluated in the MNI space. Accordingly, prior studies have implemented white matter tract templates. For example, the most predominantly used white matter tract template, the Johns Hopkins White Matter Tract Atlas, which is a white matter tract atlas based on 81 individuals (resolution: 2.5 x 2.5 x 2.5 mm) (Hua *et al*., 2008), has been used to evaluate microstructural deficits in AD (Kantarci *et al*., 2017; Araque Caballero *et al*., 2018). Here, we conducted probabilistic tractography in 100 Human Connectome Project subjects (resolution: 1.25 x 1.25 x 1.25 mm) (Van Essen *et al*., 2013). Using well-established methods to create white matter tract templates (Archer *et al*., 2018), we have provided a newly available white matter tract atlas of the uncinate fasciculus, parietal component of the inferior longitudinal fasciculus, and cingulum. In addition to recently available templates of the fornix (Brown *et al*., 2017) and tapetum (Archer *et al*., 2019), this atlas provides significantly more coverage of the brain compared to prior templates (see **Supplemental Figure 1**). Despite these strengths, this study used a cohort which is both highly educated and primarily non-Hispanic white individuals, thus limiting the generalizability to other cohorts. Further, while free-water imaging is a novel technique to quantify both extracellular and intracellular microstructure in a dMRI image, it is still unclear what cellular processes contribute to each variable. However, since our novel medial temporal lobe tract templates are freely available, we are confident that future studies can easily incorporate these into studies and begin to further elucidate these mechanisms.

In conclusion, this study provided compelling evidence that changes in hippocampal volume and FW white matter metrics in tracts projecting from the hippocampus co-occur and synergistically interact. Findings provide additional evidence that AD is a network-level disease, with white matter alterations tightly coupled with gray matter changes, and the downstream consequences of gray matter atrophy. White matter and gray matter metrics of damage are likely complementary, and both should be thoughtfully incorporated into theoretical models of aging and AD.

## Supplemental Material

**Supplemental Table 1 --.** Baseline Tract Microstructure x Hippocampal Interaction Associations

| Tract | FA<sub>T</sub> |  | FW |  |
| --- | --- | --- | --- | --- |
|  | β (95% CI) | *p*-value | β (95% CI) | *p*-value |
| **Baseline Composite Memory Performance** |  |  |  |  |
| Inferior Longitudinal Fasciculus | -6.72 (-11.1 to -2.35) | 0.36 | 2.55 (0.41 to 4.68) | 0.47 |
| Tapetum | -7.58 (-13.24 to -1.92) | 0.45 | 2.22 (0.19 to 4.26) | 0.50 |
| Uncinate Fasciculus | -2.99 (-7.77 to 1.81) | 0.71 | 0.6 (-1.34 to 2.54) | 0.91 |
| Cingulum Bundle | -8.3 (-13.31 to -3.29) | 0.36 | 1.84 (-0.53 to 4.22) | 0.67 |
| Fornix | -0.82 (-5.92 to 4.29) | 0.92 | -0.02 (-1.51 to 1.46) | 0.99 |
| **Baseline Composite Executive Function Performance** |  |  |  |  |
| Inferior Longitudinal Fasciculus | 7.67 (3.38 to 11.96) | 0.36 | -1.52 (-3.64 to 0.61) | 0.68 |
| Tapetum | 5.65 (0.11 to 11.19) | 0.51 | -3.38 (-5.34 to -1.41) | 0.36 |
| Uncinate Fasciculus | 12.85 (8.20 to 17.51) | 0.12 | -2.95 (-4.83 to -1.07) | 0.36 |
| Cingulum Bundle | 8.93 (4.01 to 13.84) | 0.36 | -0.66 (-2.99 to 1.67) | 0.91 |
| Fornix | 1.03 (-3.99 to 6.05) | 0.92 | -1.82 (-3.27 to -0.37) | 0.47 |

**Abbreviations:** FA<sub>T</sub>, free-water corrected fractional anisotropy; FW, free-water.

**Supplemental Figure 1.**
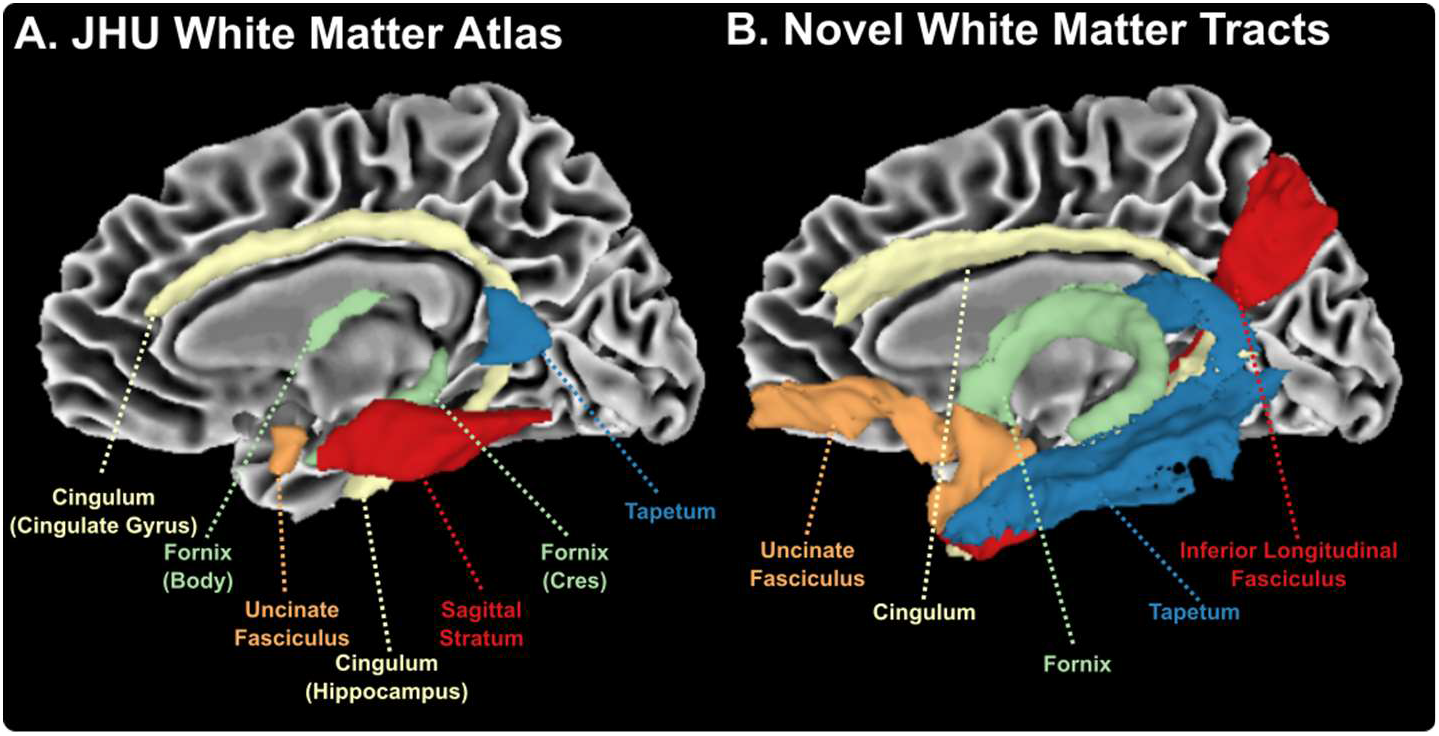
(**A**) The Johns Hopkins University (JHU) White Matter Atlas includes several medial temporal lobe projections, including the cingulum bundle, fornix, uncinate fasciculus, sagittal stratum, and tapetum. The cingulum bundle is split into two components (cingulate gyrus and hippocampus). The fornix is also splint into two components (body and cres). The sagittal stratum includes both the inferior longitudinal fasciculus and inferior fronto-occipital fasciculus. (**B**) The novel white matter tracts provided in this study (uncinate fasciculus, cingulum, inferior longitudinal fasciculus), in addition to recently available templates of the fornix and tapetum, provide more comprehensive coverage of the medial temporal lobe projections.

